# Braconidae revisited: *Bracon brevicornis* genome showcases the potential of linked-read sequencing in identifying a putative *complementary sex determiner* gene

**DOI:** 10.1101/2020.07.20.211656

**Authors:** K. B. Ferguson, B. A. Pannebakker, A. Centurión, J. van den Heuvel, R. Nieuwenhuis, F. F. M. Becker, E. Schijlen, A. Thiel, B. J. Zwaan, E. C. Verhulst

**Affiliations:** Wageningen University & Research, Laboratory of Genetics, Wageningen, The Netherlands; University of Bremen, FB02, Institute of Ecology, Population and Evolutionary Ecology Group, Bremen, Germany; Wageningen University & Research, Bioscience, Wageningen, The Netherlands; Wageningen University & Research, Laboratory of Entomology, Wageningen, The Netherlands

## Abstract

*Bracon brevicornis* is an ectoparasitoid of a wide range of larval-stage Lepidopterans, including several pests of important crops, such as the corn borer, *Ostrinia nubilalis*. It is also one of the earliest documented cases of complementary sex determination in Hymenoptera. Here, we present the linked-read genome of *B. brevicornis*, complete with an *ab initio*-derived annotation and protein comparisons with fellow braconids, *Fopius arisanus* and *Diachasma alloem*. We demonstrate the potential of linked-read assemblies in exploring regions of heterozygosity and search for structural and homology-derived evidence of the *complementary sex determiner* gene (*csd*).

## INTRODUCTION

*Bracon brevicornis* (Wesmael) is a gregarious ectoparasitoid of various Lepidoptera larvae, including many important pests, and is considered a cosmopolitan species (Temerak 1983b; Venkatesan *et al*. 2009). In the past *B. brevicornis* has been classified under the genus *Habrobracon* (Speicher and Speicher 1940), *Microbracon* (Narayanan *et al*. 1954), or classified as one species with *Habrobracon/Bracon hebetor* (Puttarudriah and Basavanna 1956), however recent research shows that *B. brevicornis* and *B. hebetor* are genetically two distinct species (Kittel and Maeto 2019). In the field, *B. brevicornis* has shown potential as a biological control agent against important pest species in stored corn stalks, such as *Ostrinia nubilalis* and *Sesamia cretica* (Kares *et al*. 2010), or against the coconut moth, *Opisinia arenosella* (Venkatesan *et al*. 2009). In the laboratory, *B. brevicornis* attacks a wide range of larval host such as *Ephesthia kuehniella, Galleria* spp., and *Spodoptera* spp. (Temerak 1983a).

Work on *B. brevicornis* has included both laboratory and semi-field set-ups to determine both its efficacy as a biological control agent as well as its suitability as a study system. There are several studies on the biology of *B. brevicornis*, e.g. on population growth potential ((Srinivasan and Chandrikamohan 2017), their host range (Temerak 1983a), interspecific competition (Venkatesan *et al*. 2009), clutch size and fitness (Villacañas de Castro and Thiel 2017), mate choice (Thiel *et al*. 2013; Thiel and Weeda 2014), diet (Temerak 1983b), and efficacy (Kares *et al*. 2010).

Within a phylogenetic perspective, *B. brevicornis* falls within the subfamily Braconinae, the largest of the cyclostome-forming braconid wasps (Chen and van Achterberg 2019). The presence of a cyclostome (round mouthpart) is a defining feature within braconid wasps, as it represents an unresolved evolutionary and systematic question: is the cyclostome a derived trait within certain branches, or an ancestral trait that has been lost in others (Chen and van Achterberg 2019)? Within the Braconinae, there have been multiple switches from ectoparasitism to endoparasitism and *vice versa*, and this combination of cyclostome and endoparasitism has been described as a “controversial topic” by braconid researchers and taxonomists (Chen and van Achterberg 2019). These systematic issues are far from being resolved, and more genomic data would be useful for future phylogenetic analyses (Chen and van Achterberg 2019). Yet, a representative genome for the Braconinae is currently lacking. As previously stated, *B. brevicornis* is an ectoparasitoid, and its position within a family that contains both types of parasitism lifestyles holds promise for further phylogenetic comparisons.

In addition, as being part of the order Hymenoptera, *B. brevicornis* has a haplodiploid sex determination system where males develop from unfertilized eggs and females develop from fertilized eggs (Cook and Crozier 1995; Heimpel and de Boer 2008). From a genetic perspective, *B. brevicornis* belongs to an interesting genus where sex determination and diploid male production have been widely studied (*B. hebetor*, Whiting and Whiting 1925, *B. brevicornis*, Speicher and Speicher 1940, *B. serinopae* Clark, Bertrand, and Smith 1963, reviewed in van Wilgenburg, Driessen, and Beukeboom 2006; *B*. spec. near *hebetor*, Holloway *et al*. 1999; and *B. variator*, A. Thiel, pers. comm.). Indeed, the first description of the complementary sex determination (CSD) mechanism was provided for *B. hebetor* (= *B. juglandis* by Whiting 1940, reviewed in Antolin *et al*. 2003), and recent work on *B. brevicornis* and polyploidy studies include diploid male fitness as well as ploidy-dependent mate choice behaviour (Thiel and Weeda 2014).

While straightforward to detect phenotypically through the formation of diploid males following inbreeding (van Wilgenburg *et al*. 2006), the molecular mechanism underlying CSD has thus far only been resolved to a small level of detail in the honeybee *Apis mellifera* (L.) (Hymenoptera: Apidae), with the identification of the *complementary sex determiner* (*csd*) gene (Beye *et al*. 2003). Heterozygosity at this gene leads to female development, while hemi-and homozygous individuals develop into haploid and diploid males respectively. Therefore, inbreeding often leads to diploid male production in species with a CSD mechanism as it increases homozygosity. *Csd* is a duplication of *feminizer* (*fem*), a *transformer* (*tra*) ortholog (Hasselmann *et al*. 2008) that is conserved across many insect orders as part of the sex determination cascade (Geuverink and Beukeboom 2014). When heterozygous, *csd* initiates the female-specific splicing of *fem*, which then autoregulates its own female-specific splicing, ultimately resulting in female development. Within the Hymenoptera, more duplications of *tra*/*fem* have been identified in species that are presumed to have CSD (Geuverink and Beukeboom 2014), but these *tra*/*fem* duplications have not been analysed for potential heterozygosity. Also, additional hymenopteran genomes are necessary to understand the evolutionary history of *tra*/*fem* duplications and identify the genes underlying CSD. However, an assembled genome is usually haploid as areas of heterozygosity are collapsed in the final stages of assembly. Yet recent advances in sequencing and analysis gave us the ability to view heterozygous regions, known as “phases” in diploid assemblies, within a genome which allow us to investigate potential *csd* regions.

Here we report on the whole-genome sequencing of a pool of females from an isolated *B. brevicornis* strain using 10X Genomics technology that relies on linked-read sequencing (10x Genomics Inc., Pleasanton, CA, USA). Due to their long history of genetic isolation during laboratory rearing, the females in this strain are assumed to have a high level of homozygosity, whereas a *csd* locus would retain its heterozygosity. The 10X Genomics technology allows for generating phased data in which allelic variants can be identified after assembly. High-molecular weight DNA is partitioned into small droplets containing a unique barcode and adapter in such a way that only a few DNA molecules are present within each droplet. Within each droplet the DNA is broken into pieces and the barcode (Gel Bead-in-Emulsion, “GEM”) is ligated to each of the DNA fragments. This resulting library can then be sequenced on an Illumina sequence platform. In the assembly step the reads originating from the same fragment are organized by barcode and put together into synthetic long-read fragments. Importantly, it is nearly impossible that two fragments with opposing allelic-variances are together in the same droplet (Weisenfeld *et al*. 2017). This technique therefore allowed us to identify potential *csd* candidates in the female-derived *B. brevicornis* genome after sequencing by studying the phased data containing the different haplotypes. Moreover, as *B. brevicornis* is a potential biological control agent of several pests, the availability of a full genome may provide effective ways to study and improve this species to grow it into an established biological control agent for Lepidopteran pests.

## METHODS

### Species description and general rearing

Individuals of *B. brevicornis* were taken from the laboratory colony L06. The colony was initiated in 2006 from naturally parasitized *O. nubilalis* larvae collected in maize fields near Leipzig, Germany. Species identification was first carried out by Matthias Schöller and Cornelis van Achterberg based on morphological characteristics (Bernd Wührer, AMW Nützlinge, pers. comm.) Since collection, parasitoids have been reared on late instar larvae of the Mediterranean flour moth, *E. kuehniella* (Thiel and Weeda 2014). The species identity of strain L06 was recently revalidated based on molecular data and is entirely separate from its congeneric *B. hebetor* (Kittel and Maeto 2019).

### DNA extraction

Immediately following emergence, 100 to 120 female wasps were flash frozen in liquid nitrogen and ground with a mortar and pestle. Genomic DNA was extracted using a protocol modified from Chang, Puryear, and Cairney (Chang *et al*. 1993). Modifications include adding 300 μL BME to extraction buffer just before use. Instead of 10M LiCl, 0.7 volume isopropanol (100%) was added to the initial supernatant, after which it was divided into 1.5 mL Eppendorf tubes as 1 mL aliquots for subsequent extractions. The initial centrifugation step occurred at a slower rate and for a longer period of time to adjust for machine availability. Final pellets were dissolved in 50 μL autoclaved MQ and recombined at the end of the extraction process (1.0 mL). DNA concentration was measured with an Invitrogen Qubit 2.0 fluorometer using the dsDNA HS Assay Kit (Thermo Fisher Scientific, Waltham, USA) with final assessments for DNA quality, amount, and fragment size confirmed via BioAnalyzer 2100 (Agilent, Santa Clara, California, USA).

### 10X Genomics library preparation and sequencing

As the genome of *B. brevicornis* is relatively small for the scale of the 10X platform, there is a higher risk of overlapping fragments within single GEMs. In order to reduce this risk, genomic DNA of a larger and previously analysed genome (Tomato, *Solanum lycopersicon* (L.) (Solanaceae), commercial variety Heinz 1607) (Hosmani *et al*. 2019) was used as ‘carrier DNA’. DNA extraction of *S. lycopersicon* followed the protocol of Hosmani *et al*. (Hosmani *et al*. 2019). The DNA of both *B. brevicornis* and *S. lycopersicon* was pooled in a 1:4 molar ratio.

One nanogram of this pooled DNA was used for 10X Genomics linked-read library preparation following the Chromium Genome Reagent Kits Version 1 User Guide (CG-00022) (10x Genomics, Pleasanton, USA). Barcoded linked read DNA fragments were recovered for final Illumina library construction (Illumina, San Diego, USA). The library was used for 2 × 150 bp paired-end sequencing on one lane of an Illumina HiSeq 2500 at the business unit Bioscience of Wageningen University and Research (Wageningen, The Netherlands). Sequencing data was then used for basecalling and subsequent demultiplexing using Longranger (v2.2.2) (10X Genomics) (command -mkfastq), yielding 212,910,509 paired-end reads with a read length of 150 bp.

### Assembly

To filter sequence data from Heinz tomato (*S. lycopersicon*) carrier DNA sequences, 23bp (16bp GEM + 7 bp spacer) were removed from forward reads and all reads were subsequently mapped to an in-house high quality reference assembly of the Heinz genome using BWA-MEM v0.7.17 (Li 2013). Using samtools v1.9 (Li *et al*. 2009), all unaligned read pairs (`-F=12`) were extracted and labelled non-Heinz. The assembly of the non-Heinz labelled read set was performed with 10X Supernova assembler v2.1.0 (10X Genomics), using default settings including commands for both pseudohap (`--style=pseudohap`) and pseudohap2 (`--style=pseudohap2`) outputs (Weisenfeld *et al*. 2017). These commands determine the output from Supernova, the first being the final scaffold output (pseudohap), while the second is the so-called ‘parallel pseudohaplotype’ (pseudohap2) scaffolds that represent areas of divergence or phases (Weisenfeld *et al*. 2017). Phasing is flattened in the pseudohap output by selecting the region with higher mapping coverage, whereas in the pseudohap2 output is differentiated by “.1” and “.2” at the end of each scaffold name to denote phasing, though not all scaffolds are phased at this point due to lack of divergence during assembly.

To verify whether there were no Heinz leftovers in the assembly, minimap2 v2.17-r941 (Li 2018) was used to align the assembly against the same Heinz assembly. Further examination on presence of possible non-*B. brevicornis* scaffolds, i.e. bacterial scaffolds from sample microbiome, was performed with BlobTools (v1.0) (Laetsch and Blaxter 2017), relying on megaBLAST against the NCBI NT-NR database (Acland *et al*. 2014)(2018-11-19) (`max_target_seqs=1, max_hsps=1, evalue=1e-25`) for taxonomical classification and BWA-MEM mapping of reads against scaffolds for coverage statistics. Reads mapping only against “Arthropoda” classified scaffolds were then extracted and used for a final k-mer analysis using jellyfish v2.1.1 `(-C m=21 -s=2000000000`) (Marçais and Kingsford 2011) and GenomeScope (Vurture *et al*. 2017) to infer heterozygosity.

Assembly completeness was determined using *BUSCO* (v3.0.2) with the `insect_odb9` ortholog set and the fly training parameter (Simão *et al*. 2015) while assembly statistics were determined using QUAST (Gurevich *et al*. 2013). The aforementioned pseudohap2 scaffolds were used in *csd* analysis, while the pseudohap scaffolds are now the assembly used for annotation.

### *Ab initio* gene finding and protein comparison

The coding sequences of two additional braconids (members of the subfamily Opiinae, and similar to the Braconinae belonging to the cyclostome subgroup (Li *et al*. 2013; Chen and van Achterberg 2019)) were used for gene prediction and protein comparisons: *Fopius arisanus* (Sonan) (Hymenoptera: Braconidae) and *Diachasma alloeum* (Muesebeck) (Hymenoptera: Braconidae). Both sets of coding sequences were retrieved from the NCBI Assembly Database, version ASM8063v1 for *F. arisanus* and version Dall2.0 for *D. alloem* (Acland *et al*. 2014; Geib *et al*. 2017; Tvedte *et al*. 2019).

For gene prediction, Augustus (v2.5.5) was first used to predict genes from the *B. brevicornis* assembly (Stanke and Morgenstern 2005). Using BLAST, coding sequences of *F. arisanus* were set as a query to the genome of *B. brevicornis* using default parameters (except `minIdentity=50`) (Camacho *et al*. 2009). The result was converted into a hints file that was used to predict the genes of *B. brevicornis* using *Nasonia vitripennis* (Walker) (Hymenoptera: Pteromalidae) as the species parameter in Augustus (`--species=nasonia – extrinsiccCfgFile=extrinsic.E.cfg`).

After prediction, the protein sequences were retrieved and compared to both *F. arisanus* and *D. alloeum* (version Dall2.0) using Proteinortho (v6.0, `-p=blastp, - e=0.001`) (Lechner *et al*. 2011). From the orthology grouping generated by Proteinortho, gene names could be allocated to the predicted genes. Lengths of both these *B. brevicornis* genes and the orthologs of *F. arisanus* and *D. alloem* were retrieved using samtools for comparison (Li *et al*. 2009). Errors within the annotation related to genome submission and validation were corrected with manual annotation of exons (three cases) and removal of two predicted genes that were more than 50% ambiguous nucleotides.

### *In silico* identification of *feminizer* as a putative *csd* locus

The pseudohap2 files were deduplicated using the dedupe tool within BBTools (sourceforge.net/projects/bbmap/) (`ac=f`) to remove all parallel pseudohaplotypes that were complete duplicates as these scaffolds were not heterozygous. The remainder of the set contained both scaffolds that previously had a duplicate, as well as solitary scaffolds that did not have a partner scaffold. These unique scaffolds were removed using the “filter by name” tool in BBTools, leaving 258 scaffolds, or 129 pairs of pseudohap2 scaffolds. Pairs were pairwise aligned in CLC Genomics Workbench 12 (Qiagen, Hilden, Germany) using default settings (`gap open cost=10, gap extension cost=1, end gap cost=free, alignment=very accurate`).

A local tBLASTn search against the entire *B. brevicornis* assembly was performed using the *Apis mellifera* Feminizer protein (NP_001128300) as query in Geneious Prime 2019.1.3 (http://www.geneious.com, (Kearse *et al*. 2012)). The protein of gene “g7607” (locus tag = BBRV_LOCUS33129) was used in an NCBI BLASTp against the nr database with default settings (Camacho *et al*. 2009; Acland *et al*. 2014). Next a region stretching from ∼10Kbp upstream and downstream of the first and last tBLASTn hit in scaffold 12, respectively, was annotated using HMM plus similar protein-based gene prediction (FGENESH+, Softberry, http://www.softberry.com/) with *Nasonia vitripennis tra* (NP_001128299) and *N. vitripennis* for the specific gene-finding parameters (Solovyev 2007). Only this combination of settings resulted in a full-length annotation from TSS to poly-A with seven exons. The resulting protein prediction was used in a BLASTp search with default settings against the nr database. To annotate the potential *fem* duplication, a stretch of ∼10Kbp directly upstream of the annotated putative fem was again annotated using FGENESH+ (Softberry) with *Nasonia vitripennis tra* (NP_001128299) and *N. vitripennis* for the specific gene-finding parameters (Solovyv et al. 2007). The predicted annotation contained five exons but lacked the last coding segment with stop codon. A protein alignment was made in Geneious Prime 2019.1.3 with *A. mellifera csd* (ABU68670) and *fem* (NP_001128300); *N. vitripennis tra* (XP_001604794) *and B. brevicornis* putative *fem* and *B. brevicornis* putative *fem* duplicate (*fem1*), using MAFFT v7.450 with the following settings: `Algorithm=auto, Scoring matrix=BLOSUM62, Gap open penalty=1.53, Offset value=0.123` (Katoh 2002; Katoh and Standley 2013).

### Microsynteny analysis

A microsynteny analysis was achieved by comparing the arrangement of a set of homologous genes directly upstream and downstream of *tra* or *fem* in *A. mellifera* and *N. vitripennis* using a combination of the online tool SimpleSynteny (Veltri *et al*. 2016) and tBLASTn searches using default settings in Geneious Prime. The scaffolds containing *fem* (*A. mellifera*, scaffold CM000059.5, 13.2Mbp in length), *tra* (*N. vitripennis*, scaffold NW_001820638.3, 3.7Mbp in length) or the putative *fem* (*B. brevicornis*, scaffold 12, 4.5 Mbp in length) were extracted from their respective genomes (*Apis*: GCA_000002195.1_Amel_4.5_genomic, *Nasonia*: nvi_ref_Nvit_2.1, *Bracon*: *B. brevicornis* assembly from this study) and searched with protein sequence from the following genes: *tra* (GeneID: 00121203), LOC100121225, LOC100678616, LOC100680007 originating from *N. vitripennis*; and *fem* (GeneID:724970), *csd* (GeneID:406074), LOC408733, LOC551408, LOC724886 originating from *A. mellifera*. The advanced settings for SimpleSynteny were as follows: `BLAST E-value Threshold=0.01, BLAST Alignment type=Gapped, Minimum Query Coverage Cutoff=1%, Circular Genome Mode=Off`. If the gene was not found within the extracted scaffold, it was searched for in the full genome assembly. For the image settings, `Gene Display Mode=Project Full-Length Gene`. This generated image was used together with results from the tBLASTn searches as template to draw the final figure. The final figure that we present in the Results and Discussion section depicts ∼0.9Mbp of genomic region for all three species.

### Data availability

Raw sequence data for *B. brevicornis* after removal of carrier DNA and contamination, as well as the assembly, can be found in the EMBL-EBI European Nucleotide Archive (ENA) under BioProject PRJEB35412, however, are currently being updated due to error in initial upload. In the meantime, both the assembly file (.fasta) (https://doi.org/10.6084/m9.figshare.12674189.v2) and the complete annotation file (.gff) (https://doi.org/10.6084/m9.figshare.12073911.v2) are available in a separate repository. Contaminated pseudohap scaffolds are available for download alongside the two pseudohap2 FASTA files, more details are provided in the supplementary materials at https://doi.org/10.17026/dans-xn6-pjm8.

## RESULTS

A total of 172 ng of *B. brevicornis* DNA was extracted, which was then reduced to 1 ng/μL for library preparation. Sequencing of the Heinz diluted library resulted in a total yield of 54 Gbp of data (corrected for 10X 23bp segment of forward reads). Mapping against the Heinz genome assembly showed a mapping percentage of 84.8%. There was a total of 30,278,915 unmapped pairs, comprising ∼8.39 Gbp of data. This corresponds to the 4:1 ratio between Heinz and *B. brevicornis* DNA in the library. Further scaffold decontamination with BlobTools resulted in a separation of the assembly into *B. brevicornis* scaffolds and microbiome scaffolds. The final genome is 123,126,787 bp (123 Mbp) in size, comprised of 353 scaffolds (5.5% ambiguous nucleotides). This is similar to the projected physical genome size of 133 Mbp (J. G. de Boer, unpublished data, flow cytometry). BUSCO analysis indicates a completeness of 98.7% (single orthologs 97.0%, duplicate orthologs 1.7%).

K-mer analysis of the *B. brevicornis-*only read set showed an expected haploid genome length of ∼115 mbp (105 Mbp unique, 10 Mbp repeat) and a heterozygosity of ∼0.54%. Peak coverage was 27x.

### *Ab initio* gene finding and protein comparison

In total, 12,686 genes were predicted, with an average coding sequence length of 529.86 amino acids. The number of genes correspond well to those found in *F. arisanus* (11,775) and *D. alloem* (13,273), the two closest relatives of *B. brevicornis* for which public data is available. Proteinortho analysis resulted in 7660 three-way orthology groups (7,830 *B. brevicornis* genes), while 362 othology groups contained proteins of *B. brevicornis* and *F. arisanus* (382 *B. brevicornis* genes), and 451 groups contained *B. brevicornis* and *D. alloem* genes (479 *B. brevicornis* genes). A large number of orthology groups (2,492) had no *B. brevicornis* genes, while 3,995 predicted genes remain ungrouped.

Compared to *F. arisanus*, the mean relative length of predicted *B. brevicornis* genes was 1.016, while the mean relative length for the two- and three-way orthology groups was 0.996. Similar results were obtained for comparisons to *D. alloem*, where mean relative length for *B. brevicornis* genes was 1.011 and 0.988 for the two- and three-way orthology groups. Furthermore, the pairwise lengths of all these proteins resemble each other very well (Figure 1).

**Figure 1.**
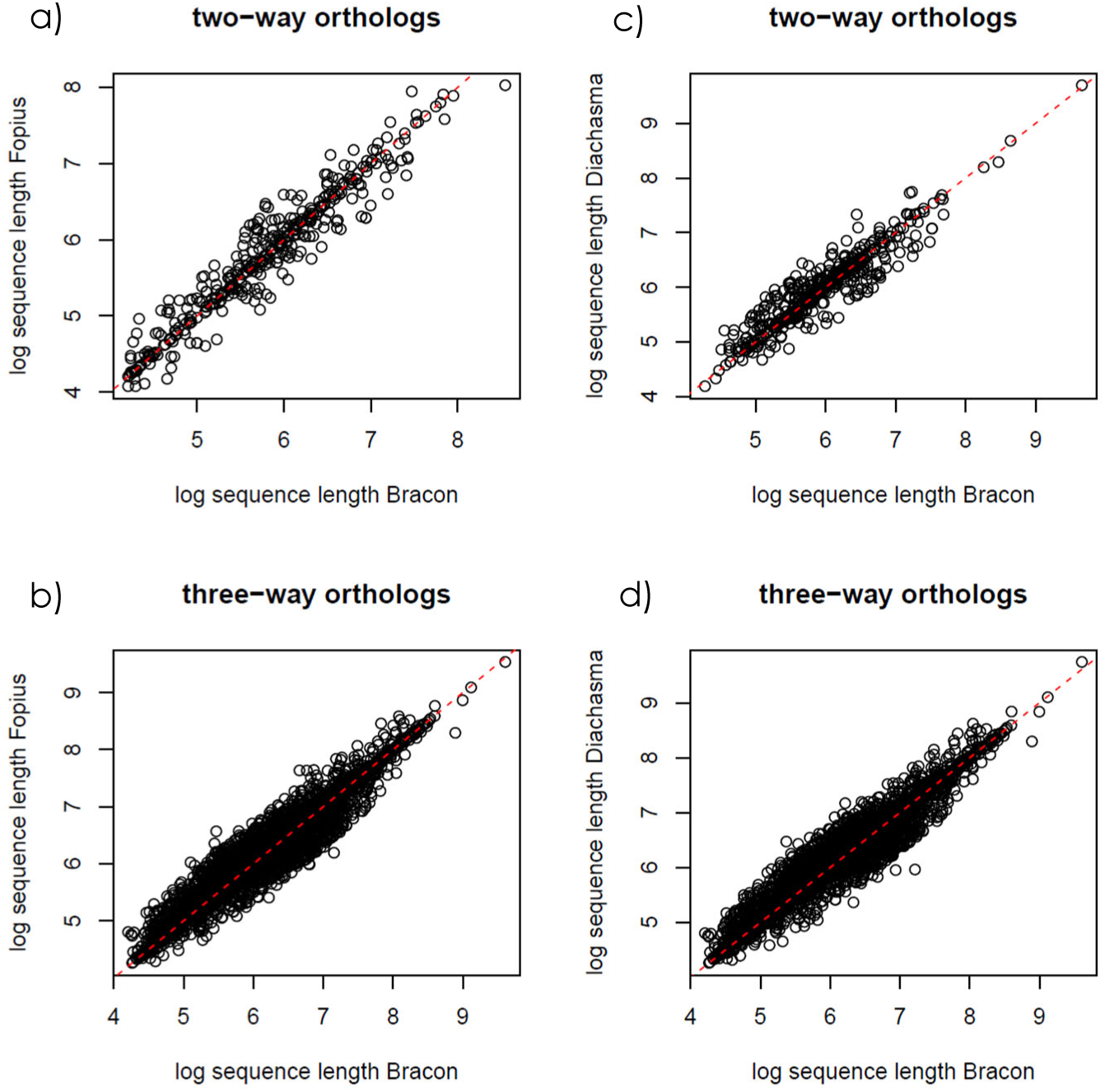
Protein length comparison between *Bracon brevicornis* and *Fopius arisanus*, a) two- and b) three-way orthologs, and *B. brevicornis* and *Diachasma alloem*, c) two- and d) three-way orthologs. Sequence lengths have been log-transformed; red dashed line indicates synteny.

### Identification of a putative *feminizer* ortholog and duplication event

After deduplicating the similar parallel pseudohaplotype files, 6,706 scaffolds in total, the remainder of the set contained 3,420 scaffolds, of which 3,286 scaffolds were solitary and did not have a counterpart pseudohap2 for comparison. Some had had a previous duplicate removed in the deduplication, while others never had a partner scaffold in the first place. These unique scaffolds were removed, leaving 258 scaffolds, or 129 pairs of pseudohap2 scaffolds. These putatively heterozygous scaffolds were good candidates to search for potential *csd* loci as these are presumed to be heterozygous in females.

So far, a *csd* gene has been sequenced only in species of bees of the genus *Apis*, and it is highly polymorphic, even within subspecies (Wang *et al*. 2012). It is located adjacent to the more conserved *feminizer* (*fem*) (Hasselmann *et al*. 2008), and we therefore started with localizing *feminizer* in the genome. As *feminizer* (or its ortholog *transformer, tra*) was not identified in the *ab-initio* annotation, we used a local tBLASTn search to find *fem* in the assembly. Four hits with E-value from 5.86e-04 to 8.59e-08 were found in scaffold 12. Searching the annotation using part of the tBLASTn result shows that it is annotated as “g7607” (locus tag = BBRV_LOCUS33129) which gave a first hit with protein O-glucosyltransferase 2 (*Diachasma alloeum*) after a BLASTp search, and no *fem* or *tra* hits were found. A closer inspection showed that “g7607” is annotated as fusion protein with the N-terminal part resembling *fem* and the C-terminal part putatively encoding *O-glucosyltransferase 2*. Next, we used FGENESH+ to re-annotate the genomic region, resulting in a full-length putative *B. brevicornis feminizer* (*Bbfem*) ortholog containing seven exons (Figure 3). We found that the two *fem*/*tra* signature domains in Hymenoptera, the Hymenoptera domain (Verhulst *et al*. 2010) and CAM domain (putative autoregulatory domain) (Hediger *et al*. 2010), are present in the putative *fem* ortholog, but are also duplicated upstream of putative *Bbfem*. A second manual re-annotation step showed that a partial *fem*-duplicate is encoded directly upstream of putative *Bbfem* containing five exons (Figure 3), which we denote here as *Bbfem1* as suggested by Koch *et al*. (Koch *et al*. 2014). The level of potential heterozygosity in the area encoding *Bbfem* and *Bbfem1* is the highest when compared across all 129 pairs of pseudohap2 scaffolds (Figure 3).

A protein alignment showed that the full-length putative *Bbfem* as well as *Bbfem1* are highly similar to each other and both contain all known *fem*/*tra* domains (Figure S1). *Bbfem1* lacks a notably long first Arginine/Serine (RS)-rich region which is present only in *Bbfem* (124-153aa), but it otherwise appears to encode for a full-length protein. The *csd-*specific hypervariable domain (Figure S1, purple text; (Beye *et al*. 2003)) is not present in *Bbfem* nor in *Bbfem1*. Therefore, the gene name has been updated as “g7607 putative Bbfem-Bbfem1 *csd*” in the official annotation.

### Synteny analysis of putative *fem* encoding region

We compared the orthologous gene arrangement of a number of genes up- and downstream of *N. vitripennis <u>tra</u>* and *A. mellifera fem* and *csd*, with the genomic organization of the *Bbfem* region (Figure 2). *N. vitripennis* LOC100680007 is present in the *tra*/*fem* containing scaffolds of all three genomes, while *A. mellifera* LOC408733 has both translocated closer to *Nasonia tra* and to a different scaffold in *B. brevicornis. N. vitripennis* LOC100121225 and LOC100678616 are encoded in opposing directions in both *A. mellifera* and *N. vitripennis* but are both downstream of *tra* in *N. vitripennis* and upstream of *fem* and *csd* in *A. mellifera*. There is no match for both genes in *B. brevicornis. A. mellifera* LOC724886 and LOC551408 are encoded in opposing directions with the same orientation in both *N. vitripennis* and *A. mellifera* but are reversed in *B. brevicornis* and downstream of *Bbfem* and *Bbfem1* while they are upstream of *csd* and *fem* in *A. mellifera*. In *N. vitripennis* both genes are not located in the *tra* containing scaffold but in another scaffold indicating that this region has undergone chromosomal rearrangements.

**Figure 2.**
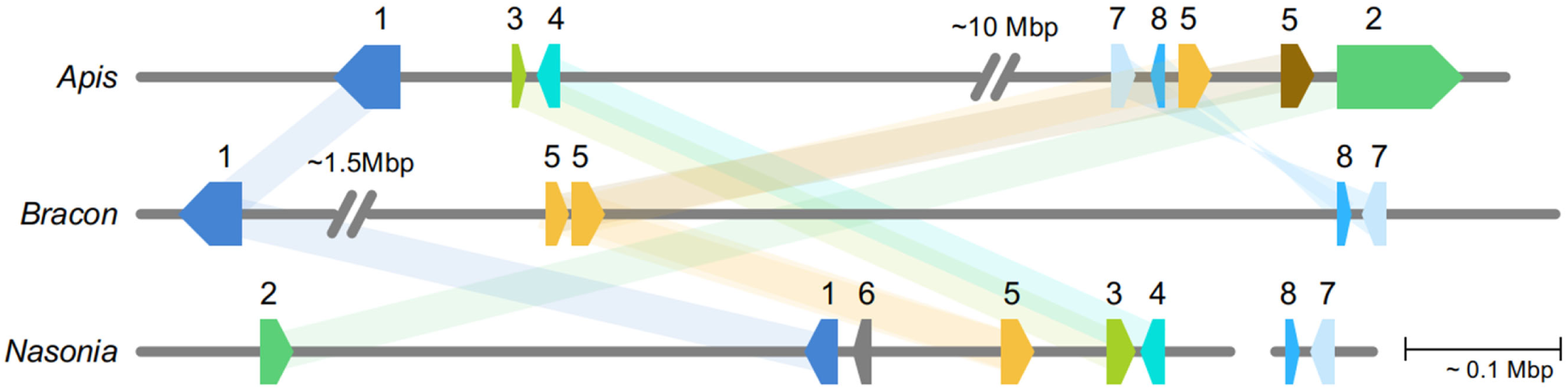
Microsynteny of genomic regions containing *tra*/*fem* paralogues. Shown is ∼0.9 Mbp of genomic region of *Apis mellifera, Bracon brevicornis* and *Nasonia vitripennis*, containing the approximate coding region for 1. LOC100680007 (dark blue), 2. LOC408733 (green), 3. LOC100121225 (lime), 4. LOC100678616 (cyan), 5. *tra*/*fem*/*fem1* (yellow) and *csd* (brown), 6. LOC107980471 (gray), 7. LOC724886 (blue), 8. LOC551408 (light blue). Locus 2 is located on a different scaffold in *B. brevicornis*, locus 3 and 4 are not present in *B. brevicornis*. Locus 6 is unique to *N. vitripennis*, and locus 7 and 8 are located on a different scaffold in *N. vitripennis*, which is depicted on the right. Both 7 and 8 are in the same order and orientation as in *B. brevicornis*, but reversed in *A. mellifera*.

**Figure 3.**
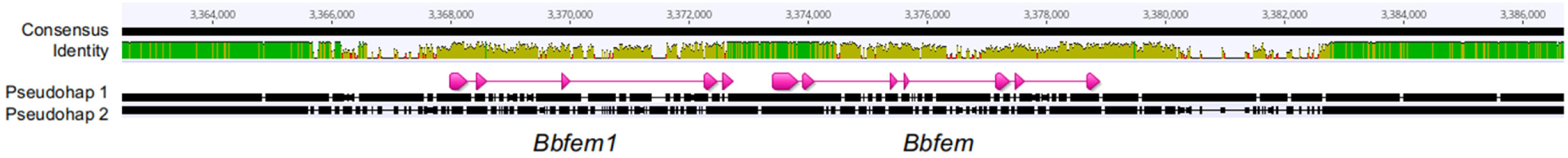
*Bracon brevicornis* annotation of *Bbfem* and *Bbfem1* on the alignment of pseudohaplotype track 1 and 2 in Geneious Prime 2019.1.3 (http://www.geneious.com, (Kearse *et al*. 2012)). Within the assembled genome, this section corresponds to a region on scaffold 12. The *Bbfem1* annotation lacks the last coding segment with stop codon. The identity track shows the amount of sequence identity across an arbitrary window (depending on zoom setting) and can be used as a proxy for heterozygosity. Green is identical, yellow is mismatch, and red is no match due to introduced gaps during alignment. The coding regions of *Bbfem1* and *Bbfem* are in a high putatively heterozygous region.

## CONCLUSIONS AND PERSPECTIVES

Here, we present the genome of the braconid wasp *Bracon brevicornis*, a parasitoid wasp that not only has biological control applications, but also offers potential as a study system for future analyses into braconid phylogenetics and gene evolution. With no previous genomes available for the subfamily Braconinae, the most specious of the braconid wasps, the resources and investigations presented here fill this gap. Our linked-read library, assisted by carrier DNA of *S. lycopersicon*, has resulted in a highly contiguous, very complete assembly, comprised of just 353 scaffolds and 12,686 genes. This gene count is similar to related species, and in further protein length comparisons, the proteins are highly similar. This indicates that the predicted genes are highly complete, a necessary feature for any future phylogenetic comparisons between species or families.

We utilized the 10X Genomics linked-read approach to obtain pseudohaploid information that would allow us to search for potential *csd* loci *in silico*. As a substantial number of scaffolds were putatively heterozygous, we used the notion that in *A. mellifera, csd* is located adjacent to *fem* (Hasselmann *et al*. 2008) to limit our search for *csd* candidates. We manually annotated a putative *B. brevicornis fem* and a partial *Bbfem* duplicate that is highly similar, and both genes encode all known *tra*/*fem* protein domains (Figure S1) (Verhulst *et al*. 2010). Both genes are in a small region that is highly heterozygous, especially when compared to the remainder of the scaffold, which would suggest true heterozygosity and not assembly error, but also when compared to the level of heterozygosity in the other 128 aligned pseudohap2 scaffolds.

Our synteny analysis showed only little structural conservation between *B. brevicornis, A. mellifera*, and *N. vitripennis* with the translocation of LOC408733 (*A. mellifora*) and the absence of LOC100121225 and LOC100678616 (*N. vitripennis*) in the *B. brevicornis* genome region. It is known that genomic regions encoding sex determination genes are dynamic in nature, showing both duplications and translocations (Dechaud *et al*. 2019). Also, *tra*/*fem* duplications have been shown in CSD systems before, most notably in *A. mellifera* where a *fem* gene duplication event resulted in it becoming a *csd* locus (Hasselmann *et al*. 2008; Gempe *et al*. 2009). However, also in non-CSD systems *tra* duplications have been observed (Geuverink and Beukeboom 2014; Jia *et al*. 2016; Geuverink *et al*. 2018). Although there is some debate on whether *fem* paralogs originated due to a single duplication event and functions as *csd* (Schmieder *et al*. 2012), or evolved multiple times independently and may have other functions (Koch *et al*. 2014), we suggest that the *Bbfem* paralog, *Bbfem1*, is a good *csd* gene candidate in *B. brevicornis*. However, in-depth analyses are required to verify this. Ultimately, our presented genome with its pseudohaploid information provides multiple opportunities for future studies, such as to improve the biological control opportunities with this species, but also to shed light on the evolutionary history of complementary sex determination systems.

## ACKNOWLEDGEMENTS AND FUNDING

We would like to acknowledge Jetske de Boer for information on *B. brevicornis* genome size, Martin Hasselmann for discussion on honeybee *csd*, and Elzemiek Geuverink for discussions on *tra*/*fem* duplicates in Hymenoptera. This project was funded by the European Union’s Horizon 2020 research and innovation program under the Marie Sklodowska-Curie grant agreement no. 641456.

## SUPPLEMENTARY MATERIALS

Additional supplementary material from this study (contaminated scaffolds, pseudohap2 scaffolds) are available on the DANS EASY Repository, https://doi.org/10.17026/dans-xn6-pjm8

**Figure S1.**
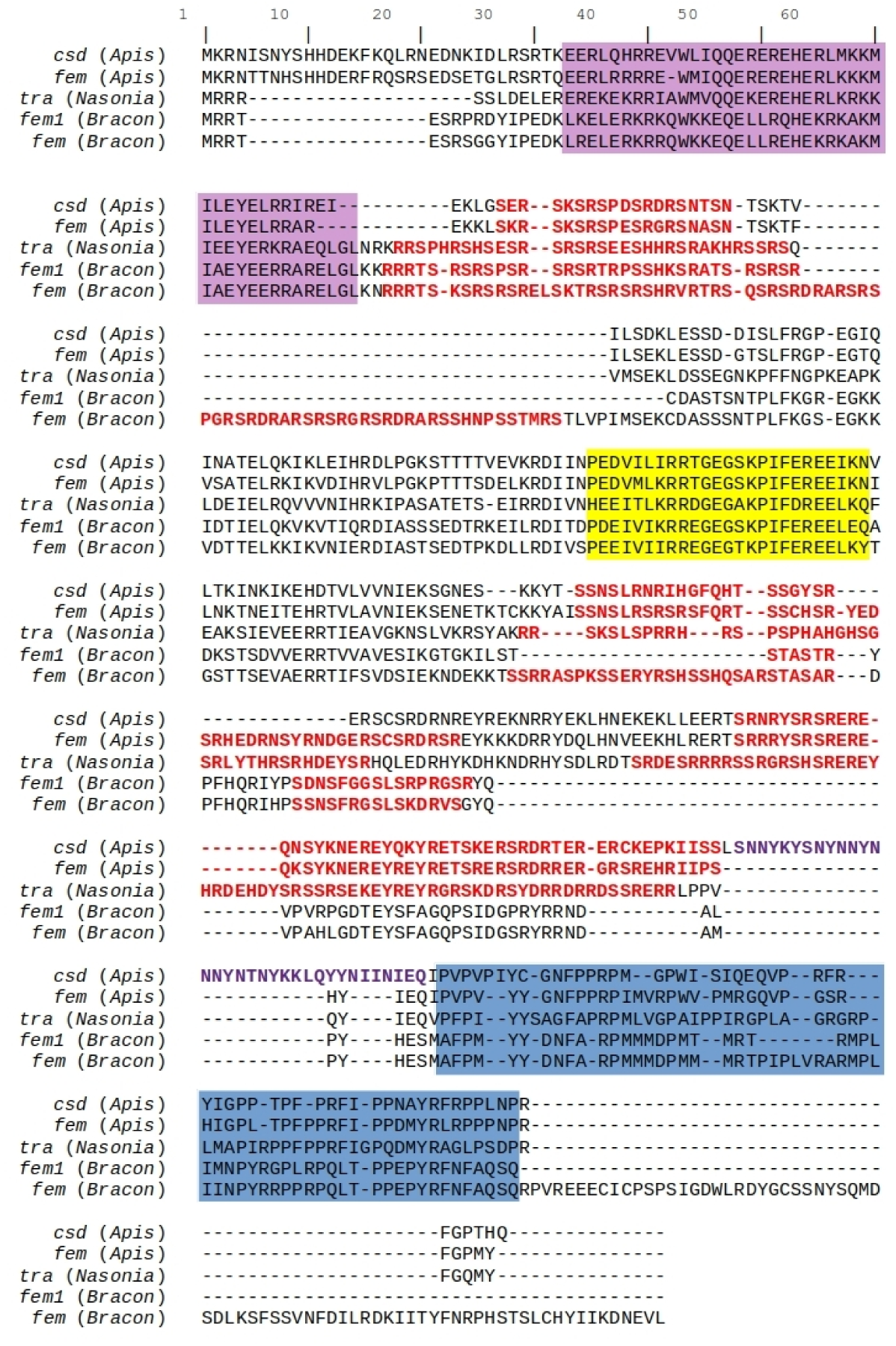
Protein alignment of *A. mellifera csd* (ABU68670) and *fem* (NP_001128300), *N. vitripennis tra* (XP_001604794), *B. brevicornis fem* and *fem1*. Purple shading indicates Hymenoptera domain (Verhulst *et al*. 2010), yellow shading indicates CAM domain (Hediger *et al*. 2010), blue shading indicates Proline (P)-rich region, red text colour indicates Arginine/Serine (RS)-rich regions, and purple text colour indicates hypervariable region in *csd* (Beye *et al*. 2003).

